# In silico model of basal ganglia Deep Brain Stimulation in Parkinson’s Disease captures range of effective parameters for pathological beta power suppression

**DOI:** 10.1101/2025.06.30.662262

**Authors:** Mahboubeh Ahmadipour, Alberto Mazzoni

## Abstract

In Parkinson’s disease (PD), the beta band ([12 30] Hz) component of basal ganglia activity is pathologically high. Deep brain stimulation (DBS) is an effective treatment to suppress symptoms of PD and is known to suppress pathological beta activity. However, the mechanism underlying this effect is not completely understood. Here, we tested the circuital effects of DBS in a computational model of the basal ganglia network in dopamine-depleted condition mimicking PD. Our model reproduces suppression of beta pathological oscillations in the basal ganglia network induced by subthalamic nucleus (STN) DBS. Crucially, this occurs for realistic levels of DBS intensity only if we incorporate short-term plasticity in STN projections to their downstream targets. STN stimulation hampers beta oscillations in the subthalamo-pallidal beta loop. This induces a progressive dephasing between this loop and the striato-pallidal beta loop, which leads in turn to a network-wide suppression of beta oscillations. This is also reflected in a restoration of the balance between D1 and D2 firing rates, which was altered by dopamine depletion. Moreover, the model also reproduces the DBS-induced gamma activity associated with symptom recovery. Finally, we explored the circuital effects of a broad range of DBS parameters in suppressing beta oscillations, focusing on the clinically relevant range of 60-150 Hz stimulation. The model suggests that 60-80 Hz stimulation frequencies might achieve beta desynchronization for an intensity even lower than the one needed at the standard 130 Hz frequency. Overall, our model lays the ground for in-silico tests of a broad spectrum of stimulation patterns.

**Author Summary:** Deep brain stimulation (DBS) is an effective treatment for alleviating symptoms of Parkinson’s disease (PD), but its underlying mechanism is unclear. Motor symptoms of PD are linked to exaggerated beta-band oscillations in a particular brain area called basal ganglia (BG). In this study, we developed a computational model to simulate how DBS affects brain circuits involved in PD. We found that stimulation can reduce pathological beta rhythms at realistic intensities only when we include realistic synaptic dynamics that temporarily adjust connection strength based on recent activity. Our model shows that stimulation decouples two available beta loops in the BG network and helps restore normal patterns of activity. Interestingly, we also found that stimulation at 60-80 Hz might be effective at lower intensities than the standard 130 Hz. Our work provides new insights into how DBS functions and supports safe and efficient exploration of a wide range of stimulation patterns.

## Introduction

Parkinson’s disease is the second most common neurodegenerative disorder, predominantly recognized for its motor symptoms, including tremor at rest, rigidity, bradykinesia, akinesia, and postural instability [1]. Beyond motor impairments, PD also encompasses a spectrum of non-motor symptoms, such as sleep disturbances, constipation, mood disorders, autonomic dysfunction, and fatigue [2]. Motor symptoms are associated with loss of dopaminergic neurons in the substantia nigra pars compacta (SNc) and consequent alterations in neural dynamics of basal ganglia (BG) [3–5]. Dopamine depletion destroys the balance in striatal activity by increasing the firing rate of dopamine-inhibited D2 neurons and decreasing the firing rate of dopamine-excited D1 neurons [4], with pathological beta oscillations [12–30 Hz] subsequently emerging in the striatum (STR), subthalamic nucleus (STN), globus pallidus pars externa (GPe), and globus pallidus pars interna (GPi) [6–11]. Several studies investigated the generation mechanism through which dopamine depletion leads to these pathological oscillations [12–23]. The first hypothesis suggests that beta oscillations emerge from the interaction between GPe and STN [12–14], while other studies propose that the cortex [15–17] or striatum [18] may be the origin. Another hypothesis emphasizes the significant contribution of the interaction between GPe and the striatum in driving these oscillations [19–22]. In our previous study, we suggested that the synchronization of two loops (STN-GPe loop and GPe-striatum loop) that are independently oscillating in the beta range is the key factor responsible for these abnormal oscillations [23]. For high levels of dopamine, the two loops are decoupled, and the oscillation power is low while with dopamine depletion the two loops synchronize and produce the pathological oscillations.

Deep brain stimulation is a neuromodulation technique which alleviates motor symptoms of PD, but its mechanism is not clear yet. One key hypothesis is local suppression, where somatic firing is reduced via mechanisms like GABAergic activation, synaptic depression, or depolarization blockade [24–26]. Several studies investigated the downstream, and their findings were contrary to what was expected according to suppression of target area since STN-DBS increased pallidal firing [27] despite the excitatory STN-pallidal projections and pallidal DBS reduced thalamic firing [28] despite the inhibitory pallidal-thalamus connections. Therefore, it was proposed that while somatic firing may be suppressed, efferent axons and synaptic terminals are activated at stimulation frequency, leading to the axon-soma decoupling hypothesis [29,30]. An alternative explanation is that DBS activates the afferent axons converging on target neurons. According to this explanation, the distribution of afferent inputs (whether mainly inhibitory or excitatory) and the local neuroanatomical features would determine whether DBS produces a net local inhibitory or excitatory response [31]. Additionally, it was hypothesized that DBS induces antidromic (toward cortex) action potential propagation [25,32]. In addition to modulating average firing rate, DBS significantly alters the oscillatory dynamics in PD: Numerous studies have shown that high-frequency stimulation reduces exaggerated beta-band oscillations, which are closely linked to motor impairments in PD [33–37]. DBS has also been shown to enhance gamma band activity, which is often associated with improved motor performance [33,38]. Gamma oscillations are not unique to DBS; they have been consistently observed across basal ganglia structures in PD patients with dopaminergic treatment [39–41], where they are dynamically modulated by movements [42,43]. However, elevated gamma activity has been linked to the development of dyskinesias in some patients, highlighting its dual role in Parkinson’s disease [44].

To better understand the mechanisms underlying DBS effects, computational models have been developed to simulate basal ganglia circuits under healthy, Parkinsonian, and DBS conditions [45–54], ranging from detailed spiking neuronal networks to mean-field approaches. One notable limitation of most current modeling studies is their omission of short-term plasticity (STP), a key mechanism that can significantly impact network behaviour and signal processing. STP, which refers to transient changes in synaptic strength over milliseconds to minutes that influence the integration and timing of neuronal signals across network, plays a critical role in shaping neural circuit dynamics [55]. Here, we hypothesized that STP plays a crucial role in the context of DBS because DBS activates efferent axons and synaptic terminals at the stimulation frequency, leading to repeated synaptic activity. However, the release of neurotransmitters and the resulting post-synaptic currents are influenced by the synapse’s firing history [56], which reflects STP mechanisms such as synaptic facilitation and depression that dynamically regulate synaptic strength based on recent activity patterns. In this study, we extended our previously established computational basal ganglia model [23] by incorporating the biophysical Hanson and Jaeger synaptic plasticity [57] to investigate the impact of STN-DBS on basal ganglia dynamics under Parkinsonian conditions. We modelled the effect of STN-DBS as activation of efferent axons close to DBS electrode. Our simulations showed that STN-DBS suppresses beta oscillations in BG and enhances gamma oscillations in striatal medium spiny neurons. In addition, we investigated how STN-DBS can suppress beta activity and the effect of stimulation parameters on the beta power. The results revealed that STN DBS desynchronizes the two principal loops which are responsible for the generation of beta oscillations.

## Results

### Interplay between Stimulation Intensity and Synaptic Plasticity in DBS-induced beta power suppression

We investigated whether a simple model of STN DBS effects on a spiking network model of dopamine-depleted BG (see Methods and Fig 1A) is able to replicate the neural activity modulations induced by DBS on STN observed in clinics. It is well known that PD related dopamine depletion leads to aberrant beta power in the STN, and this effect is captured by our model (compare orange and green line in Fig 1B). The most relevant effect of DBS on STN is the decrease in beta range power as a function of the stimulation intensity. Indeed, in our model, progressively increasing the number of neurons activated by DBS (see Methods) results in the restoration of levels of beta power similar to the ones of the non-dopamine depleted network (Fig 1B). Stimulating approximately 40% of STN neurons was necessary to suppress beta activity to normal levels, aligning with clinical observations that indicate activation of an adequate volume of neurons is essential for achieving therapeutic benefits [70,71]. However, when synaptic plasticity was not included, the model produced an unrealistic outcome: stimulating only 8% of STN neurons was sufficient to eliminate excessive beta power and restore normal levels. This contradicts experimental and clinical findings, where DBS typically requires a broader stimulation of the STN neurons to effectively reduce pathological symptoms [72,73]. This discrepancy suggests that synaptic plasticity within the basal ganglia network plays a crucial role in shaping the effects of DBS and must be accounted for to develop realistic models of PD.

**Fig 1.**
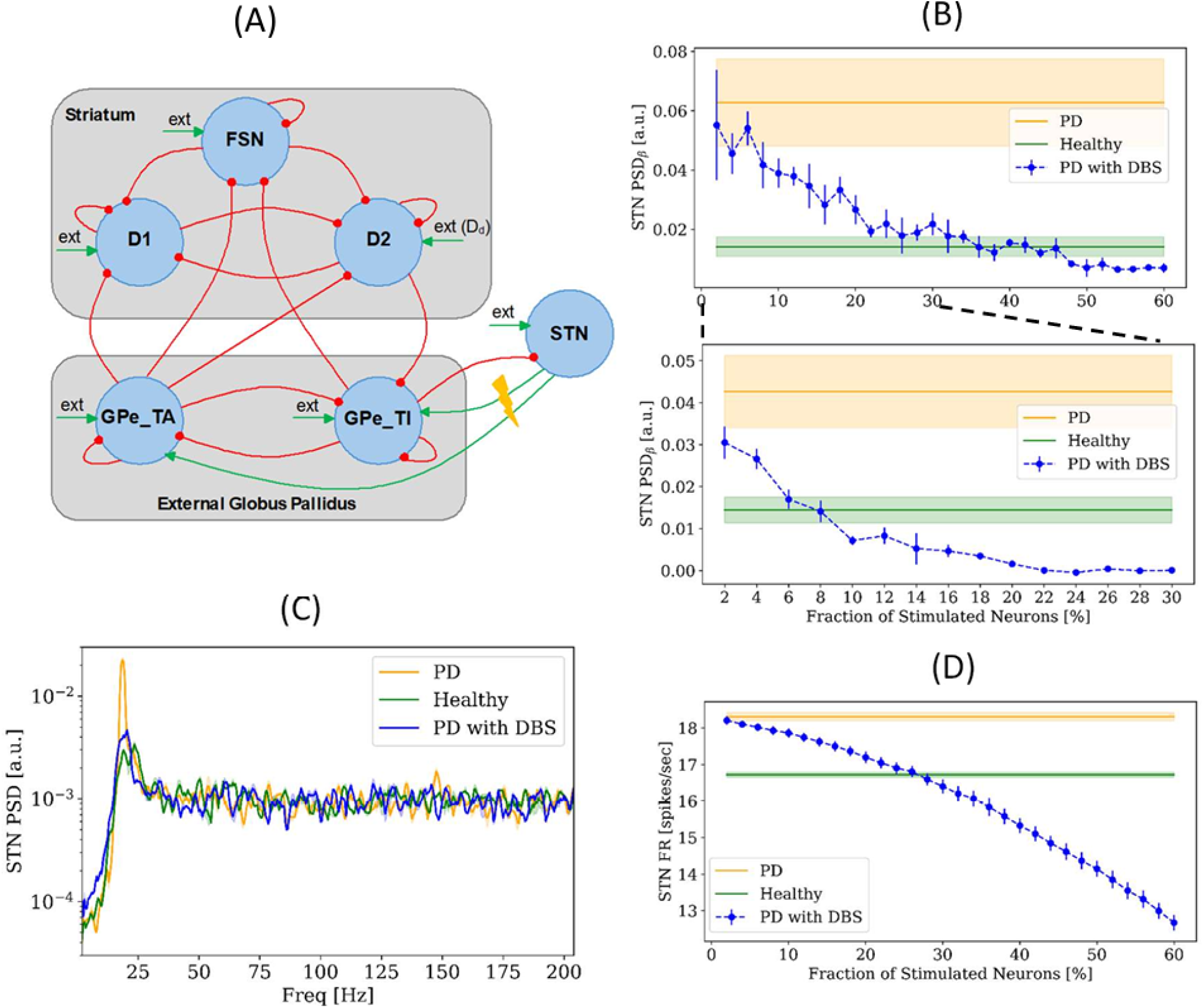
Effect of STN-DBS on STN. (A) Architecture of the spiking BG network model with DBS applied to the STN. FSN, D1 and D2: striatal fast spiking neurons, and medium spiny neurons with D1 and D2 dopamine receptors; GPe-TA and GPe-TI: arkypallidal and prototypic populations of the globus pallidus pars externa; STN: subthalamic nucleus; ext: external excitatory Poissonian input. Red/green arrows indicate inhibitory/excitatory projections. Dopamine depletion (D_d_, external drive) was modelled by adjusting the excitatory Poissonian input to D2 neurons. DBS effect was modelled by replacement of projection rate of a fraction of STN neurons to GPe. (B) Variation of STN beta power in relation to fraction of stimulated neurons with (top) and without (bottom) plasticity. STN beta power in healthy (green) and parkinsonian (orange) conditions are also presented. (C) Power spectral density (PSD) of STN in healthy (green), parkinsonian (orange) and parkinsonian with STN-DBS at 40% intensity (blue) conditions in BG network model with plasticity. (D) Variation of STN firing rate in relation to fraction of stimulated neurons in BG network model with plasticity.

Considering the stimulation of 40% of neurons as an effective DBS condition, we compared the power spectral densities across all conditions, finding no other significant alterations in the overall STN spectrum (Fig 1C). Increasing the fraction of stimulated neurons decreased the STN firing rate (Fig 1D) which is in line with a group of experimental results that report the inhibitory effect of DBS in target area [74–76].

### DBS Modulates Neural Activity Spectrum Across Basal Ganglia: Beyond Beta Suppression

Building on the findings from the STN-DBS effect on STN, we next explored its network-level impact on the other basal ganglia nuclei. While these regions, like STN, showed reduced beta power, the effects of STN-DBS on them extend also over other frequency ranges (Fig 2). GPe-TI and GPe-TA directly receive DBS pulses from STN axons and displayed sharp peaks at 143.5 Hz, shifted from the applied 130 Hz stimulation frequency together with the reduction of beta power 73.1% and 70.8%, respectively (Fig 2A-B). This shift is likely due to nonlinear interactions between the DBS pulses and synaptic plasticity within the BG network, which alters the oscillatory frequency induced in the GPe neurons. STN-DBS decreased beta activity in D1 and D2 while it enhanced their gamma activity (Fig 2C-D). In the absence of STN-DBS, gamma band activity was absent in D1 neurons, whereas D2 neurons exhibited gamma oscillations under Parkinsonian conditions even prior to stimulation. In D2, the decrease in beta (78.9%) was associated to an increase in the gamma range 133.7% (Fig 2D). Finally, the behaviour of FSN was similar to the one observed in STN (Fig 2E). STN-DBS altered the firing rates of these structures in parkinsonian state (Fig 2F). In the external globus pallidus, both GPe-TI and GPe-TA neurons exhibited increased activity, with firing rates rising by 17.9% and 43.2%, respectively. STN-DBS resulted in varied modulation of neuronal subtypes within the striatum. Firing rates of D1 and D2 neurons increased by 183.3% and 59.2%, respectively, while FSN exhibited 42.3% reduction in activity. STN-DBS restored the balance in striatal medium spiny neurons by increasing the firing rate of D1 neurons relative to D2 neurons, thereby resembling the healthy state.

**Fig 2.**
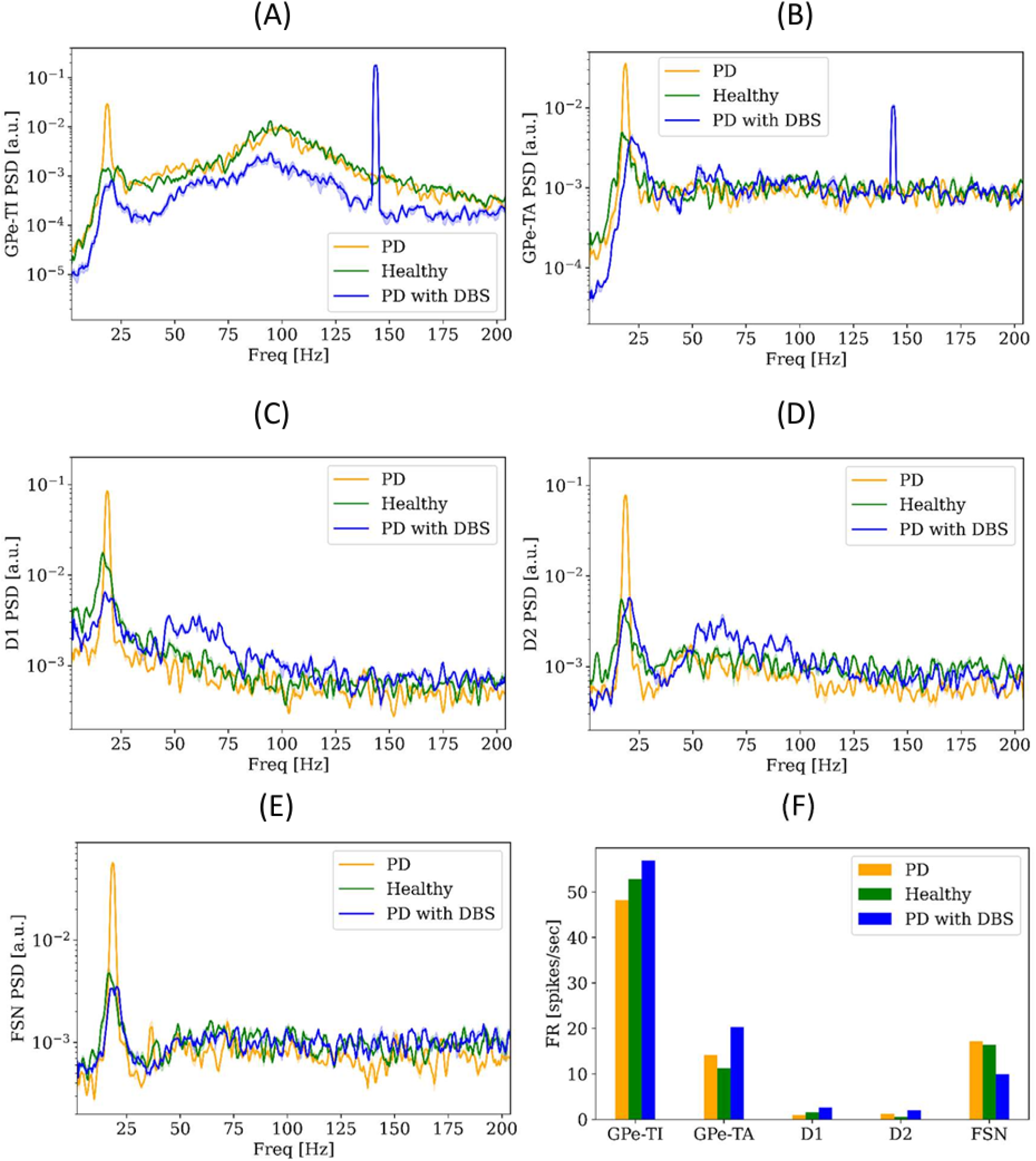
STN-DBS-induced spectral changes across nuclei. Power Spectral Densities (PSDs) of GPe-TI (A), GPe-TA (B), D1 (C), D2 (D) and FSN (E) in healthy (green), parkinsonian (orange) and parkinsonian with STN-DBS at 40% intensity (blue) conditions. (F) Firing rate of BG nuclei in healthy, parkinsonian and parkinsonian with STN-DBS at 40% intensity conditions.

### DBS Desynchronizes the Two Beta Oscillators in Basal Ganglia

To better understand how DBS suppresses pathological beta oscillations, we focused on the two principal beta-generating loops in the basal ganglia. Our previous study identified the STN–GPe-TI loop and the FSN–D2–GPe-TI loop as independent beta oscillators under healthy dopamine levels, which progressively synchronize as dopamine is depleted, contributing to the emergence of pathological beta oscillations in Parkinson’s disease [23]. Our findings in the previous sections indicated that beta power in the STN decreased as the stimulation intensity increased (Fig 1B). To understand the DBS effect on the GPe-striatum loop, we studied the effect of STN-DBS on the beta power of D2. Similar to our findings for STN, beta power in D2 decreases as stimulation intensity increases (Fig 3A). When approximately 40% of STN neurons are stimulated, D2 beta power approaches the beta power level in the healthy condition. Mean beta frequency of STN did not change by DBS while it increased for D2 as the stimulation intensity increased (Fig 3B). At around the stimulation of 40% of STN neurons, the mean beta frequencies of STN and D2 have similar values while the spectogram of the STN and D2 at 40% stimulation of STN neurons showed that the two oscillators were not synchronized (Fig 3C). To quantify the synchronization of the two oscillators in different conditions, the PLV was computed, and the results demonstrated that at high levels of dopamine (healthy condition) the two oscillators were highly independent (*PLV*≅0.3) while at low levels of dopamine (PD condition), they were highly synchronized (*PLV*≅0.7) (Fig 3D). As DBS intensity increased, the synchronization of the two oscillators decreased and at around 40% stimulation of STN neurons, the PLV reached the PLV value in healthy condition (Fig 3D).

**Fig 3.**
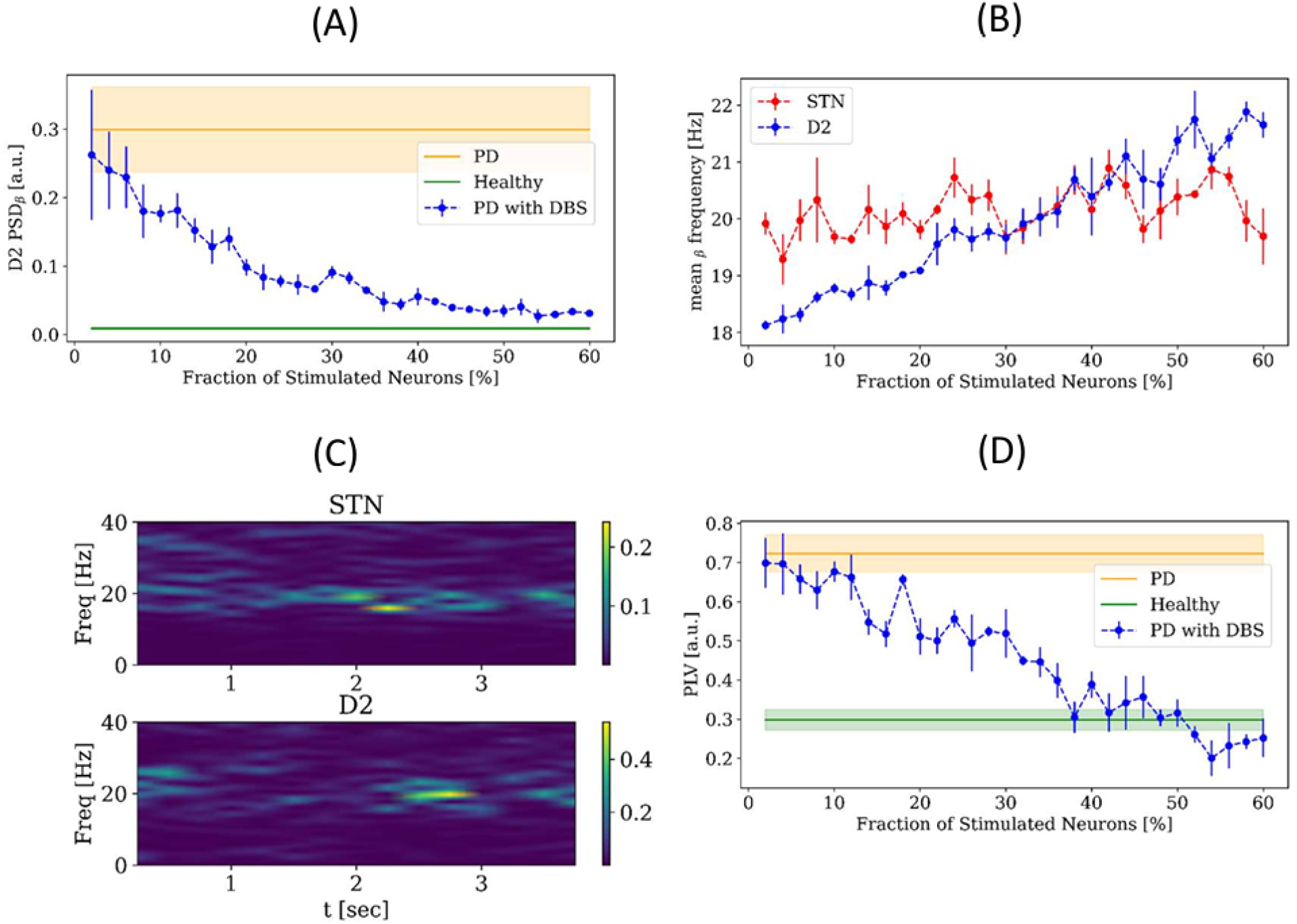
DBS-induced desynchronization of beta oscillators in basal ganglia. (A) D2 beta power in healthy (green), Parkinsonian (orange), and Parkinsonian with STN-DBS (blue), with the blue curve also showing its variation with the fraction of stimulated STN neurons. (B) Mean beta frequencies of STN (blue) and D2 (red) in relation to the fraction of stimulated neurons in STN. (C) Spectogram of STN (top) and D2 (bottom) in parkinsonian with STN-DBS at 40% intensity. (D) PLV between STN and D2 in healthy (green), Parkinsonian (orange), and Parkinsonian with STN-DBS (blue), with the blue curve also showing PLV variation with the fraction of stimulated STN neurons.

### Impact of DBS Settings on Beta Power Modulation

The results in the previous sections showed that the model can capture the main acute effects of the conventional DBS with 130 Hz frequency on basal ganglia activity. For further exploration, we studied the impact of DBS settings on DBS outcome. First, we studied the influence of DBS temporal structure on the modulation of beta power by considering Poissonian inputs as DBS pulses. Our simulations showed that delivering DBS as a Poisson process with an average frequency of 130 Hz was not able to suppress beta power in STN (Fig 4A). Indeed, the STN beta power increased as the Poissonian DBS intensity increased. This result suggests that to effectively disrupt the strong pathological beta rhythm and to restore normal BG network dynamics, a regular input is necessary. In contrast, the Poissonian input acts as white noise and is not effective in destroying the beta rhythm. Then, we investigated the impact of stimulation parameters including intensity and frequency on pathological oscillatory activity of STN (Fig 4B). Stimulation frequency was varied from 60 Hz to 150 Hz in 5 Hz increments, while intensity was defined as the fraction of STN neurons receiving stimulation, ranging from 2% to 60% in 2% steps. A heatmap was generated to visualize STN beta power across this parameter space and a white boundary line was overlaid on the plot to indicate the therapeutic threshold-the minimum stimulation intensity required at each frequency to reduce beta power to within the healthy range. Stimulation intensities above this boundary maintained beta power at or below the healthy level, indicating effective and stable therapeutic control. Interestingly, the therapeutic boundary reveals a non-monotonic pattern: stimulation frequencies in the range of approximately 85-125 Hz require relatively higher stimulation intensities to reduce beta power to the healthy level. In contrast, both lower frequencies (below ∼80 Hz) and higher frequencies (above ∼130 Hz) are associated with reduced intensity requirements. This suggests that frequencies outside the mid-range (85-125 Hz) may be more efficient in suppressing pathological beta activity, potentially reflecting complex interactions between stimulation dynamics and BG network across the frequency spectrum. This analysis provides insight into the combined role of frequency and intensity in achieving optimal DBS outcomes, supporting the use of high-frequency stimulation for Parkinsonian symptom relief, while also aligning with clinical observations [77] suggesting potential benefits of lower frequency stimulation (60–80 Hz).

**Fig 4.**
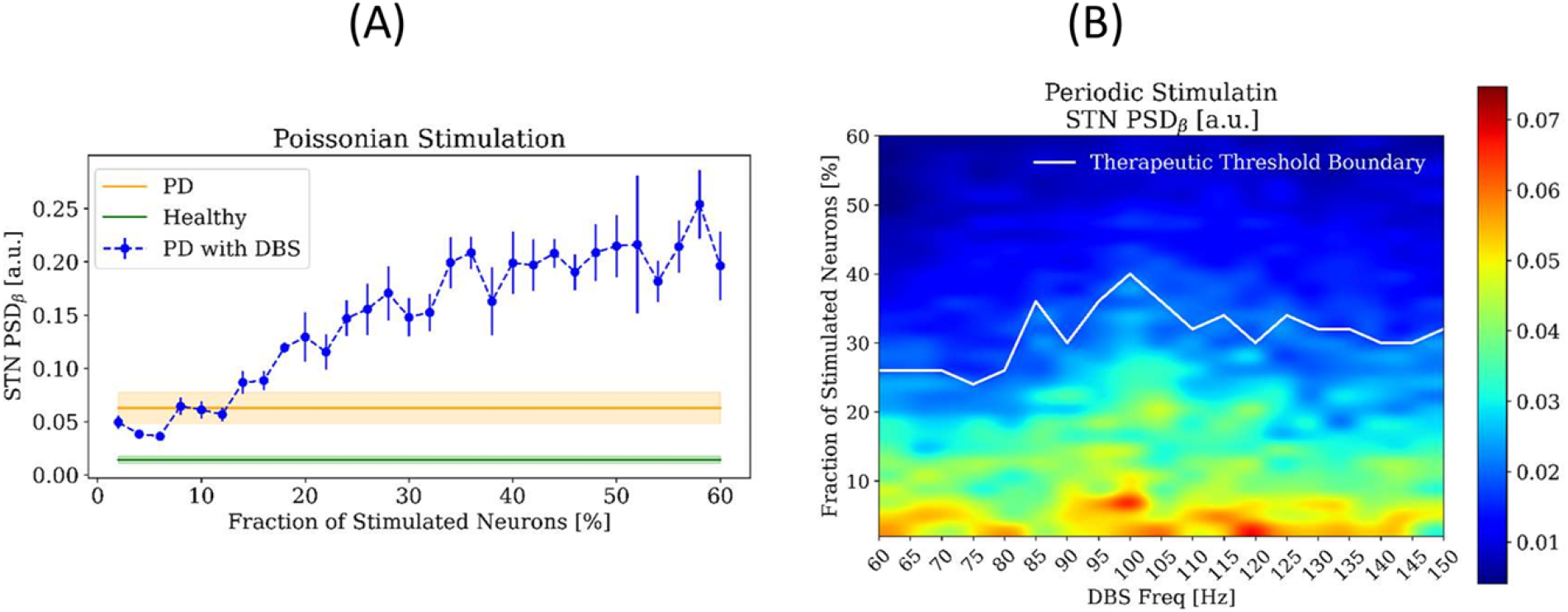
Effects of DBS settings on STN beta power. (A) STN beta power in healthy (green), Parkinsonian (orange), and Parkinsonian with Poissonian STN-DBS (blue), with the blue curve also showing its variation with the fraction of stimulated STN neurons. (B) STN beta power in relation to DBS frequency and intensity. The white boundary line indicates the therapeutic threshold, representing the minimum stimulation intensity needed at each frequency to bring beta power within the healthy range.

## Discussion

In this study, we presented a computational model of the effects of STN-DBS on the basal ganglia network. The model reproduces DBS-induced beta suppression and highlights the role of dephasing between the two beta loops in the process. Moreover, we found that STN-DBS enhances gamma band activity in the striatum, particularly in D1 and D2 receptor-type medium spiny neurons. These results are consistent with previous experimental studies showing that DBS reduces beta oscillations [33–37] and enhances gamma activity [33,38] in the basal ganglia.

We found that beta suppression is achieved for realistic ranges of intensity only if STP is taken into account. The observed beta suppression in our model is dependent on stimulation intensity, with a threshold of approximately 40% of STN neurons required to be stimulated to achieve normalization of beta power. This finding is consistent with clinical observations that sufficient stimulation volume is necessary to attain therapeutic effects [70,71]. Instead, if we do not include STP in our model, beta suppression is achieved with an unrealistic minimal stimulation. According to experimental data, high-frequency stimulation suppresses STN [24–26] and causes high-frequency projections to go downstream through axon terminals [27,28]. Given that DBS affects axonal terminals through repetitive high-frequency activation, accurately capturing synaptic dynamics is essential. Short-term synaptic plasticity, which reflects transient changes in synaptic strength based on recent activity [55,56], is particularly relevant under such conditions. To enhance the physiological realism of our model, we incorporated STP at these projections to capture the dynamic modulation of synaptic efficacy during repetitive stimulation. To simulate DBS, we modelled high frequency stimulation by replacing the physiological projections from the STN to the GPe with periodic high-frequency pulses, mimicking clinical stimulation patterns.

Previous studies primarily focused on how DBS influences thalamic relay fidelity and beta oscillations within the BG network [45–54]. The thalamus is of particular interest because motor impairments in Parkinson’s disease are believed to arise from its disability to relay sensorimotor input to the cortex. In the parkinsonian state, abnormal projection from the output nucleus of BG neuronal network (i.e., GPi) to thalamus are thought to disrupt normal thalamic relay function, leading to impaired motor control. In contrast to much of the modeling work that has centred on thalamic dynamics, Adam et al. [54] investigated the effects of STN-DBS on BG, showing that it can restore gamma band oscillations in the striatum, which are disrupted in the Parkinsonian state. Although these models have provided valuable insights into the effects of DBS, they did not account for short-term synaptic plasticity, leaving a gap in understanding how dynamic synaptic mechanisms contribute to the network’s response to DBS. Taking this into account and modeling DBS by manipulation of projections from STN to GPe showed increased activation of the GPe, particularly the GPe-TI population. Increased activity of GPe in turn exerted stronger inhibition onto the STN, recapitulating the suppressive feedback observed experimentally. However, the effects of DBS in our model extended beyond this local circuit. Neuronal populations that receive direct or indirect input from the STN, including GPe-TA and striatal neurons (D1, D2, FSNs), also exhibited significant changes in firing rates and spectral properties. Notably, STN-DBS restored the balance of striatal medium spiny neurons activity by increasing the activity of D1 neurons relative to D2 neurons, thereby approximating the firing pattern observed under healthy conditions.

In our previous study [23], we examined the circuit-level mechanisms underlying beta oscillations in the BG using a computational model. Specifically, we found that the loop between the STN and GPe-TI, as well as the loop involving striatal fast-spiking interneurons, D2 medium spiny neurons, and the GPe-TI, exhibit resonances in the beta range. While these loops function largely independently under healthy dopamine levels, their synchronization under dopamine depletion leads to an amplification of beta-band oscillations [23]. In this study, we investigated the impact of DBS on these oscillations. Our results revealed that STN-DBS impacts the synchronization of the two beta loops. STN-DBS desynchronizes these loops, with the PLV dropping to levels comparable to the healthy state when stimulation intensity reaches 40%. This supports the hypothesis that pathological synchronization of independent beta oscillators contributes to the emergence of abnormal beta rhythms in PD [23] and that DBS restores normal BG function by disrupting this synchronization. Our findings highlight the broad influence of DBS across the basal ganglia and underscore the importance of studying BG network as integrated system rather than isolated components.

One of the main objectives of the work was to provide a model enabling in silico testing of DBS parameters. Here we performed a frequency–intensity analysis revealed that achieving therapeutic effects varies non-monotonically across the 60–150 Hz range, with lower stimulation intensities required at both low (<80 Hz) and high (>130 Hz) gamma frequencies. In contrast, the mid-frequency range (∼85–125 Hz), which includes the gamma peak of GPe-TI (∼100 Hz), required higher stimulation intensities, suggesting possible interference effects that may reduce therapeutic efficiency within this band. Interestingly, although the standard clinical practice is to deliver DBS at 130-150 Hz frequency, recent results show that also 60 Hz stimulation can be clinically effective [78,79].

Our analysis of stimulation parameters further revealed that regular, structured stimulation is critical for effective suppression of beta oscillations. Poissonian DBS, which lacks temporal regularity, failed to reduce beta activity and in fact exacerbated it. This observation reinforces clinical preferences for high-frequency, regular DBS patterns and suggests that temporal structure plays a key role in disrupting pathological synchrony.

Despite the strong alignment of our results with experimental studies, our computational framework has certain limitations. Although the model incorporates key BG structures and short-term synaptic plasticity, it simplifies other aspects such as heterogeneous neuronal subtypes, and external inputs. In particular, inputs to all BG populations were modelled as uncorrelated Poisson processes, which do not fully capture their characteristics. Experimental evidence indicates that beta and gamma oscillations in the BG may, in part, be inherited from cortical activity [80–82]. Future studies could enhance biological plausibility by integrating long-term plasticity, detailed cortical and thalamic inputs, and more complex neuronal dynamics to better capture the contributions of these structures to BG function. Another limitation is that we represented dopamine loss by increasing the firing rates of D2 neurons population, capturing the enhanced activity of the indirect pathway characteristic of Parkinson’s disease. While this method successfully reproduces key pathological features, it simplifies the broader impact of dopamine depletion by excluding other important factors such as alterations in synaptic strength, plasticity mechanisms, and network-level adaptations [59]. By focusing on the primary consequence of dopamine loss, the imbalance between the direct and indirect pathways of the basal ganglia, we were able to replicate major aspects of BG dynamics in PD. Finally, it is important to remember that experimental data regarding beta oscillations consist primarily of local field potential recordings, while here we compute population outputs based on their firing rates. Future extensions of the model should aim to incorporate these additional elements, including a proper model of the relationship between firing activity and local field potential in the basal ganglia, for a more comprehensive representation of the disease state.

In summary, our findings highlight the importance of stimulation intensity and synaptic plasticity in determining the efficacy of STN-DBS. By disrupting pathological synchronization and restoring physiological oscillatory activity, DBS exerts widespread modulatory effects across the basal ganglia. These results emphasize the necessity of incorporating biologically realistic synaptic mechanisms in computational models to enhance our understanding of neuromodulation therapies and to guide the optimization of DBS protocols for Parkinson’s disease.

## Materials and methods

### Basal Ganglia model

We used a spiking neuronal network model of the basal ganglia, originally developed in [23] and subsequently updated in [58], which we further modified in this study to incorporate synaptic plasticity mechanisms. The BG network is a comprehensive biophysical model including striatum, globus pallidus externa (GPe), subthalamic nucleus (STN) and their connections (Fig 1A). Striatum consists of D1-type dopamine receptor medium spiny neurons (D1-MSN), D2-type dopamine receptor medium spiny neurons (D2-MSN) and a population of fast spiking (inter-)neurons (FSN). GPe is divided into two populations that are labelled as GPe-TA (characterized by a lower discharge rate and by a negligible input from striatal populations) and GPe-TI (with a higher activity and receiving input from D2). Each population has a specific size reported in Table 1. Neurons are connected randomly, with fixed connection probabilities between populations, based on previous studies [23,59]. Each neuron receives independent excitatory Poissonian input, simulating afferent signals from other brain regions not explicitly modelled. External input rates were adjusted across populations to align their mean firing rates with ranges reported in experimental observations: FSN [10-20] Hz ([60,61]), D1 and D2 (MSN) [0.5–2.5] Hz ([62]), GPe-TI [30–60] Hz ([20,21]), GPe-TA [10–20] Hz ([21]) and STN [12–20] Hz ([63]).

**Table 1.** Size of populations in BG network.

| Population | Number of Neurons |
| --- | --- |
| D1 | 6000 |
| D2 | 6000 |
| FSN | 420 |
| GPe-TA | 264 |
| GPe-TI | 780 |
| STN | 408 |

Neurons are modelled as conductance based, adaptive and point neurons. Adaptive exponential neurons are applied for the STN and GPe populations and their dynamics are as:

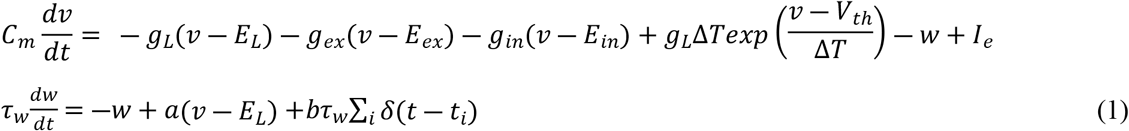

Adaptive quadratic neurons are used for striatal populations; For D1 and D2 populations, the equations are as:

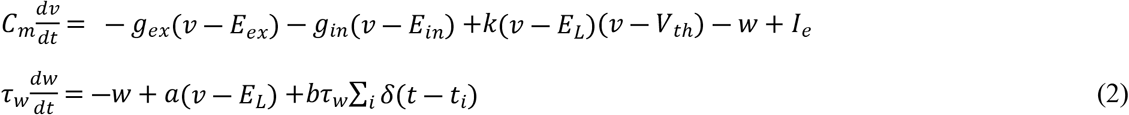

while for FSN, the dynamics are governed by:

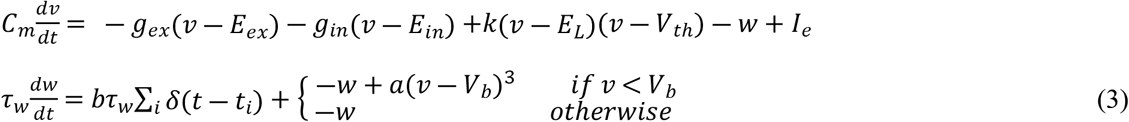

For all types of neurons, if the membrane potential reaches *V_peak_*, it will reset to *V_reset_*:

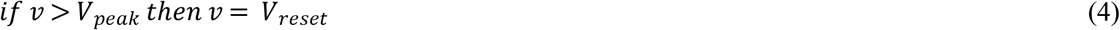

In the previous work, all the synapses were considered static while in this work we used frequency-dependent synapses to capture short-term plasticity (STP) for STN to GPe-TI and STN to GPe-TA projections. The change in conductance of static synapses presents an exponential decay:

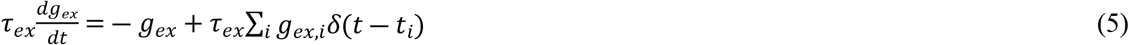

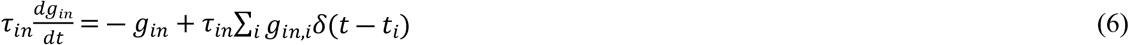

For dynamic synapses, we used the model proposed by Hanson and Jaeger that can fit physiological STP data [57]. The idea of the model is to describe facilitation and depression phenomena in synapses. The facilitation mechanism enhances the synaptic strength, and it could reflect the increase in neurotransmitters release due to the calcium influx caused by spikes arriving in the presynaptic terminal [64,65]. The depression mechanism diminishes the synaptic strength due to the consumption of neurotransmitters for the transmission of action potentials. The effective synaptic strength is determined by the combination of both facilitation and depression mechanisms.

The Hanson and Jaeger model uses two variables to describe these phenomena; variables *F* and *D* which represent facilitation and depression, respectively. The dynamics of the synapses with STP is governed by:

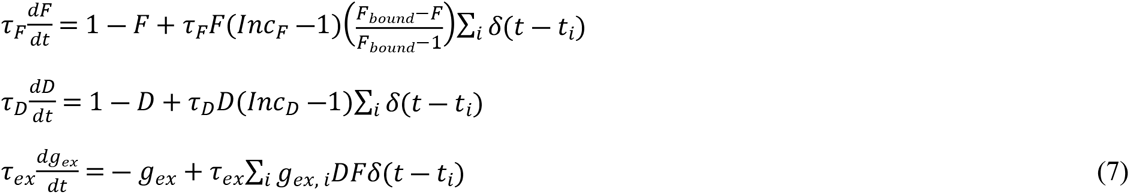

According to physiological recordings [57], each STN to GPe synapse was designated as one of the three types of plasticity, facilitation-dominant, depression-dominant or pseudo-linear with the same probability.

Parameter values for each neuronal population are reported in Table 2. The connectivity properties (delays, connection probabilities and synaptic weights) are presented in Table 3. These values were acquired from our previous studies [23,58] which has been originated from the work of Lindahl and Kotaleski [59]. The parameter values of dynamic synapses, which were extracted from Hanson and Jaeger study [57], are listed in Table 4.

**Table 2.**
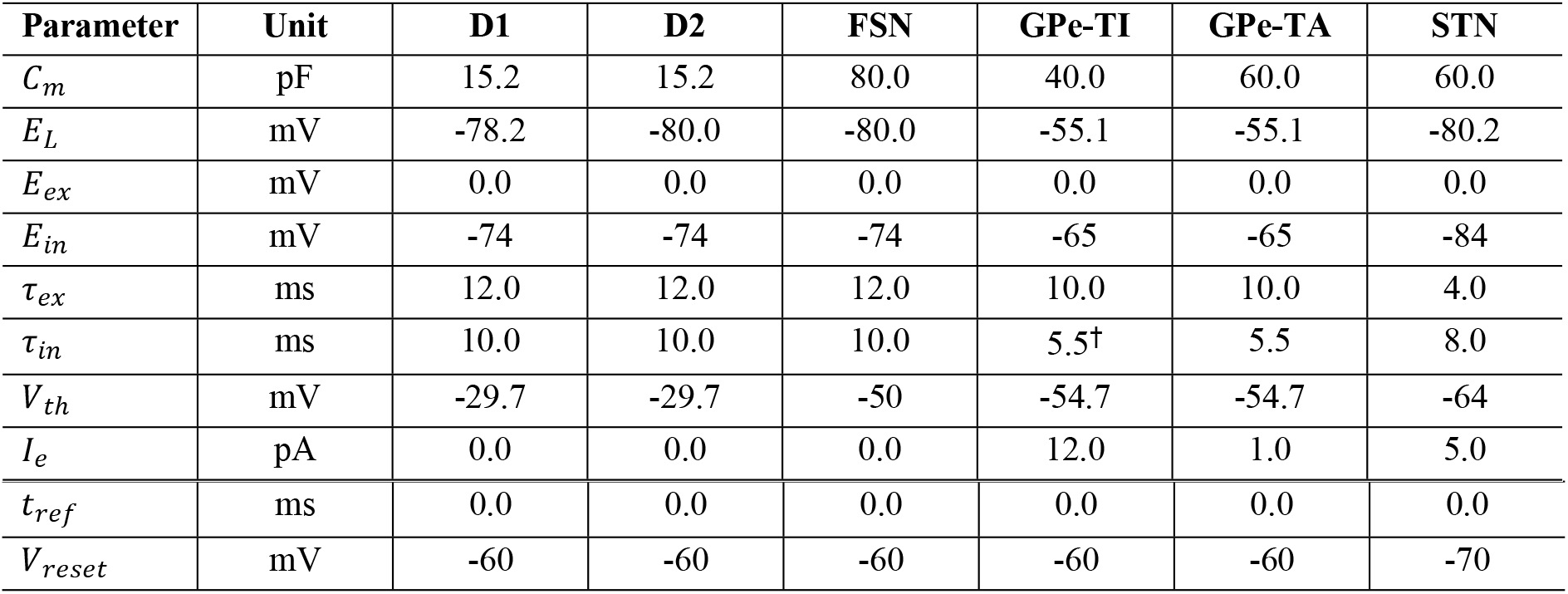
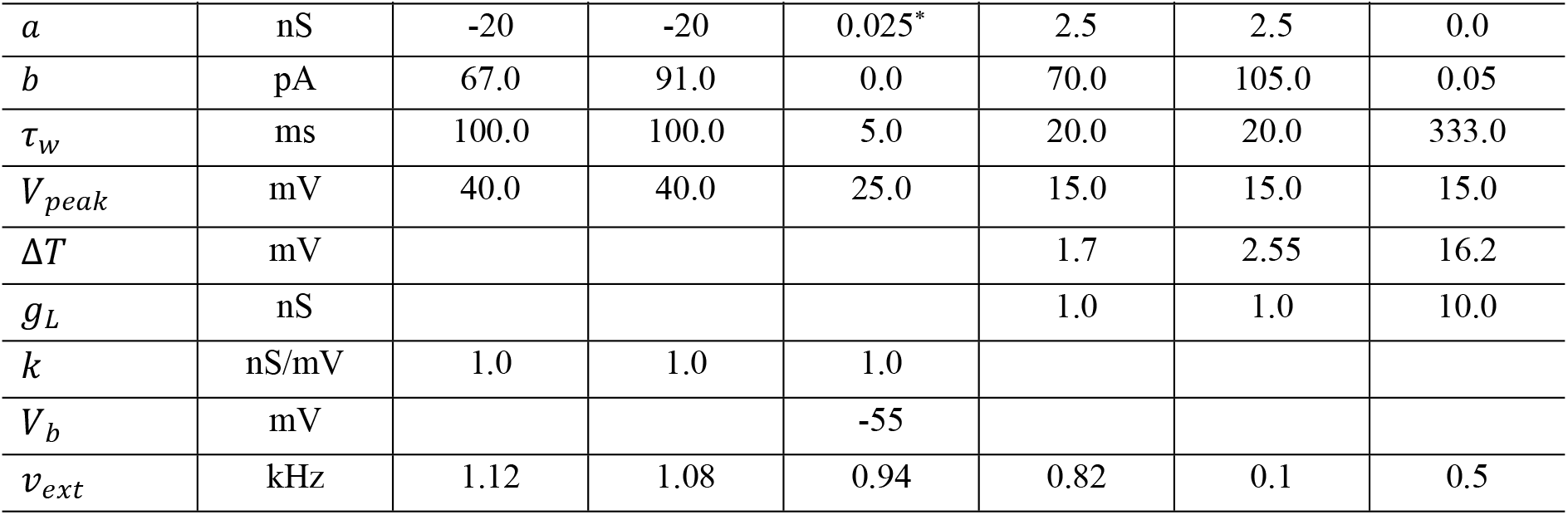
Parameters of each neuronal population. (^*^): for the FSN population the unit of a is ns/mV_2_. (†): recurrent inhibitory synapses for GPe-TI were modeled separately using a distinct conductance variable with its own time constant τ*_rec_* = 7.

| Parameter | Unit | D1 | D2 | FSN | GPe-TI | GPe-TA | STN |
| --- | --- | --- | --- | --- | --- | --- | --- |
| $C_m$ | pF | 15.2 | 15.2 | 80.0 | 40.0 | 60.0 | 60.0 |
| $E_L$ | mV | -78.2 | -80.0 | -80.0 | -55.1 | -55.1 | -80.2 |
| $E_{ex}$ | mV | 0.0 | 0.0 | 0.0 | 0.0 | 0.0 | 0.0 |
| $E_{in}$ | mV | -74 | -74 | -74 | -65 | -65 | -84 |
| $\tau_{ex}$ | ms | 12.0 | 12.0 | 12.0 | 10.0 | 10.0 | 4.0 |
| $\tau_{in}$ | ms | 10.0 | 10.0 | 10.0 | 5.5† | 5.5 | 8.0 |
| $V_{th}$ | mV | -29.7 | -29.7 | -50 | -54.7 | -54.7 | -64 |
| $I_e$ | pA | 0.0 | 0.0 | 0.0 | 12.0 | 1.0 | 5.0 |
| $t_{ref}$ | ms | 0.0 | 0.0 | 0.0 | 0.0 | 0.0 | 0.0 |
| $V_{reset}$ | mV | -60 | -60 | -60 | -60 | -60 | -70 |

**Table 2 (continued). Parameters of each neuronal population.**
| Parameter | Unit | D1 | D2 | FSN | GPe-TI | GPe-TA | STN |
| --- | --- | --- | --- | --- | --- | --- | --- |
| $a$ | nS | -20 | -20 | 0.025* | 2.5 | 2.5 | 0.0 |
| $b$ | pA | 67.0 | 91.0 | 0.0 | 70.0 | 105.0 | 0.05 |
| $\tau_w$ | ms | 100.0 | 100.0 | 5.0 | 20.0 | 20.0 | 333.0 |
| $V_{peak}$ | mV | 40.0 | 40.0 | 25.0 | 15.0 | 15.0 | 15.0 |
| $\Delta T$ | mV | | | | 1.7 | 2.55 | 16.2 |
| $g_L$ | nS | | | | 1.0 | 1.0 | 10.0 |
| $k$ | nS/mV | 1.0 | 1.0 | 1.0 | | | |
| $V_b$ | mV | | | -55 | | | |
| $\nu_{ext}$ | kHz | 1.12 | 1.08 | 0.94 | 0.82 | 0.1 | 0.5 |

**Table 3.**
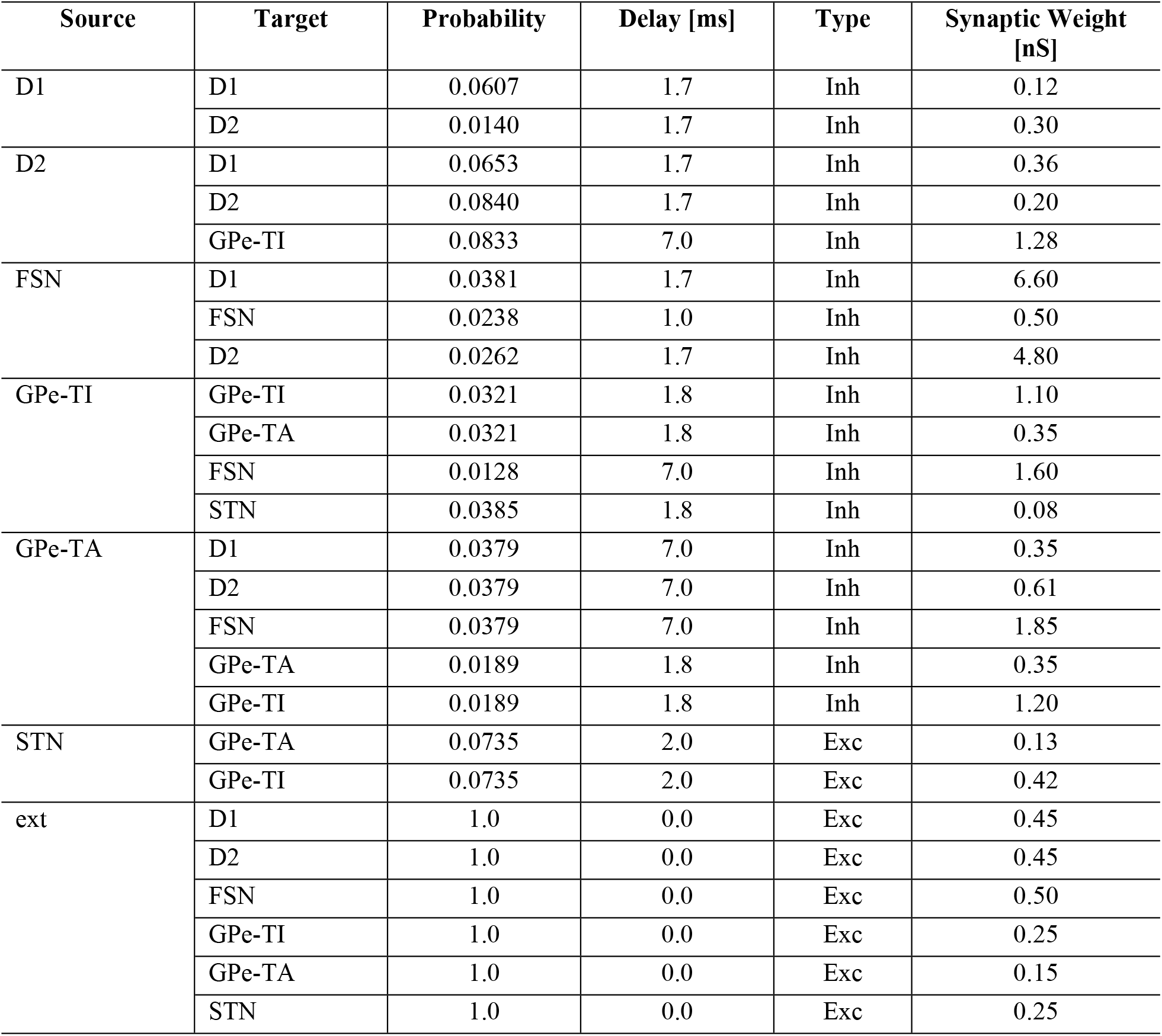
Connectivity properties of the BG model.

| Source | Target | Probability | Delay [ms] | Type | Synaptic Weight [nS] |
| --- | --- | --- | --- | --- | --- |
| D1 | D1 | 0.0607 | 1.7 | Inh | 0.12 |
|  | D2 | 0.0140 | 1.7 | Inh | 0.30 |
| D2 | D1 | 0.0653 | 1.7 | Inh | 0.36 |
|  | D2 | 0.0840 | 1.7 | Inh | 0.20 |
|  | GPe-TI | 0.0833 | 7.0 | Inh | 1.28 |
| FSN | D1 | 0.0381 | 1.7 | Inh | 6.60 |
|  | FSN | 0.0238 | 1.0 | Inh | 0.50 |
|  | D2 | 0.0262 | 1.7 | Inh | 4.80 |
| GPe-TI | GPe-TI | 0.0321 | 1.8 | Inh | 1.10 |
|  | GPe-TA | 0.0321 | 1.8 | Inh | 0.35 |
|  | FSN | 0.0128 | 7.0 | Inh | 1.60 |
|  | STN | 0.0385 | 1.8 | Inh | 0.08 |
| GPe-TA | D1 | 0.0379 | 7.0 | Inh | 0.35 |
|  | D2 | 0.0379 | 7.0 | Inh | 0.61 |
|  | FSN | 0.0379 | 7.0 | Inh | 1.85 |
|  | GPe-TA | 0.0189 | 1.8 | Inh | 0.35 |
|  | GPe-TI | 0.0189 | 1.8 | Inh | 1.20 |
| STN | GPe-TA | 0.0735 | 2.0 | Exc | 0.13 |
|  | GPe-TI | 0.0735 | 2.0 | Exc | 0.42 |
| ext | D1 | 1.0 | 0.0 | Exc | 0.45 |
|  | D2 | 1.0 | 0.0 | Exc | 0.45 |
|  | FSN | 1.0 | 0.0 | Exc | 0.50 |
|  | GPe-TI | 1.0 | 0.0 | Exc | 0.25 |
|  | GPe-TA | 1.0 | 0.0 | Exc | 0.15 |
|  | STN | 1.0 | 0.0 | Exc | 0.25 |

**Table 4.** Parameter values for STP Synapses.

| Synapse Type | $\tau_F$(ms) | $\tau_D$(ms) | $Inc_F$ | $Inc_D$ | $F_{bound}$ |
| --- | --- | --- | --- | --- | --- |
| Facilitation-dominant | 241 | 491 | 1.4 | 0.9 | 5 |
| Depression-dominant | 148 | 764 | 1.64 | 0.55 | 5 |
| Pseudo-Linear | 345 | 700 | 1.34 | 0.86 | 5 |

### Modeling of dopamine depletion

Dopaminergic degeneration in Parkinson’s disease significantly disrupts striatal function by altering the activity of medium spiny neurons (MSNs), which constitute most striatal neurons and are key components of the basal ganglia circuitry. Experimental studies have demonstrated that dopamine depletion alters the excitability of projection neurons in a receptor-specific manner, leading to suppressed activity in D1 neurons and enhanced activity in D2 neurons [66,67]. To simulate parkinsonian condition, we directly implemented the increase in D2 neuron activity by modulating the rate of its external excitatory input. Furthermore, due to the inhibitory nature of projections from D2 to D1 neurons, increased D2 activity resulted in a decreased firing rate of D1 neurons. The intensity of the external input rate towards the D2 population was modelled as:

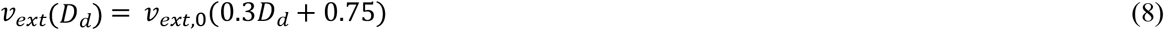

where *v_ext,0_* is the reference value of input rate (see Table 2) and *D_d_* is the parameter regulating the severity of the condition of dopamine depletion. The higher is *D_d_*, the more is dopamine depletion. Specifically, *D_d_* = 0.166and *D_d_* = 0.5 were used for simulating the behaviour of the BG in healthy and parkinsonian condition, respectively.

### Modeling of DBS effect

In the present work, we assumed that STN-DBS directly affects only a subset of STN neurons, with the proportion of affected neurons increasing with stimulation intensity. To simulate this effect, the spike trains propagating along the axons of these neurons were replaced with artificial spike trains at the DBS frequency. Let *N* be the number of STN neurons. For a given stimulation intensity, a fraction (α) of STN neurons, were considered to be stimulated and their axonal outputs were overridden by DBS spike trains *t_DBS_*, while the remaining axons preserve their natural spike times *t_STN_*.

This modeling approach is motivated by experimental studies investigated the impact of DBS on neuronal behavior. One group of studies explored the target area in which DBS was injected but used different approaches to cope with stimulus artifact; Some of them reported a reduction in neuronal activity [24] and some an increased activity of neurons in the target area [68]. A second group of studies examined downstream structures and found that DBS activates projections from target area [27]. These findings support the hypothesis that DBS exerts both inhibitory and excitatory effect [29]; That is, it could hyperpolarize cell body while still excite an action potential at the axon initial segment.

### Spectral and synchronization analysis

The activity of each population was considered as its instantaneous firing rate computed over time bins of one millisecond. Welch method (windows = 2000ms, overlap = 50%) was used to compute the power spectral density (PSD) of the activity.

To quantify the intensity of oscillations in a specific frequency band [*f*_1_ *f*_2_], for population *i* within the BG network, we computed the mean spectral power as:

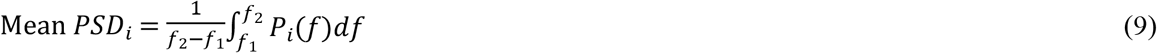

where *P_i_*(*f*) is the PSD of the nucleus activity. Specifically, we used [12 30] Hz and [30 150] Hz as the frequency bands for beta and gamma oscillations respectively. This measurement is biased due to the presence of constant activity (no oscillations). To eliminate this bias, we considered the corrected quantity as:

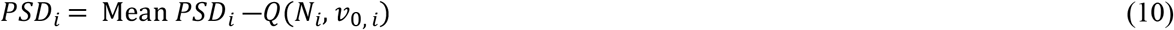

where *Q*(*N_i_*, *v*_0,_ *_i_*) is the spectral power of an auxiliary population with the same number of neurons and constant mean activity in the nucleus (*i*.*e*, *N_i_*, *v*_0,_ *_i_*). The fictitious activity of this auxiliary population has been estimated through the binomial distribution. In other words, the number of spiking neurons in each bin 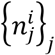 for the auxiliary population is given by independent extractions from a binomial distribution:

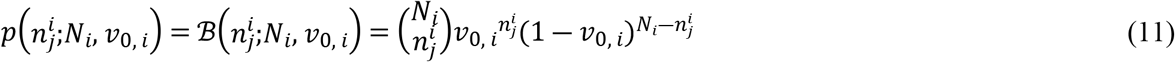

The synchronization of STN-GPe loop and GPe-striatum loop is responsible for beta exaggeration [23]. We used two approaches to measure the synchronization between the two loops in different conditions of BG. The first one is comparing the mean frequency of beta oscillations in D2 and STN as representatives of the two loops:

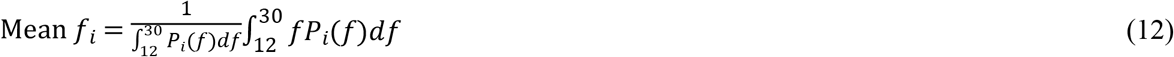

The second approach is to use phase locked variable (PLV) between the activities of STN and D2 as a measurement of the intensity of synchronization. PLV is defined as:

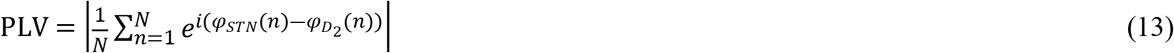

where φ*_STN_*(.) and 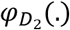 represent the phase time series of the beta range activities of STN and D2, respectively. The phase time series are computed as:

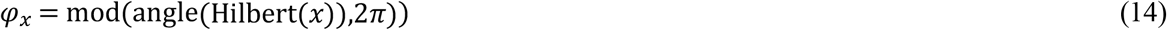

where *x* ∈ {*STN*, *D*2} represents the beta range activities of STN and D2 and is computed by bandpass filtering of STN and D2 activities with cut off frequencies of 11 and 31 Hz.

### Numerical methods

The code for simulating the BG network has been developed in Python using ANNarchy [69]. Post analysis of the simulation results has been implemented in Python as well. The fourth-order Runge-Kutta approach with a fixed time step h = 0.04 ms was used to numerically integrate the equations that describe the development of each neuron in the simulated network. The duration of each simulation was fixed to 6000 ms and the first 2000 ms was eliminated as of the transition state. For each case, 4 simulations were performed to estimate the standard deviation of computed quantities.

## Acknowledgements

The authors were supported by the Italian Ministry of University and Research (MUR), under the complementary actions to the NRRP “Fit4MedRob—Fit for Medical Robotics” Grant (# PNC0000007). The work was also supported by #NEXTGENERATIONEU (NGEU) and funded by the Ministry of University and Research (MUR), National Recovery and Resilience Plan (NRRP), project EBRAINSItaly (IR0000011) - European Brain ReseArch INfrastructureS-Italy (DN. 101 16.06.2022).

## Data availability statement

The codes necessary to reproduce the data that support the findings of this study are openly available at https://github.com/MahboubehAhmadipour/BGNetwork_STP_DBS.

## Author Contributions

Mahboubeh Ahmadipour: Conceptualization, Data curation, Formal analysis, Investigation, Methodology, Software, Validation, Visualization, Writing– original draft, Writing– review & editing; Alberto Mazzoni: Conceptualization, Investigation, Methodology, Validation, Supervision, Project administration, Funding acquisition, Writing– review & editing.

